# Modulation of corneal tissue mechanics influences epithelial cell phenotype

**DOI:** 10.1101/362236

**Authors:** Ricardo M. Gouveia, Guillaume Lepert, Suneel Gupta, Rajiv R. Mohan, Carl Paterson, Che J. Connon

**Affiliations:** Institute of Genetic Medicine, Newcastle University, International Centre for Life, Newcastle-upon-Tyne, NE1 3BZ, UK; Imperial College London, Blackett Laboratory, London, SW7 2BW, UK; Harry S. Truman Memorial Veterans Hospital, Columbia, MO, USA; College of Veterinary Medicine, University of Missouri, Columbia, MO, USA

## Abstract

Whilst the control of stem cell differentiation using substrates of differing compliance has been extensively explored *in vitro*, the significance of this mechanism at a physiological level is not known. Here we set to explore the role of corneal surface biomechanics in controlling epithelial cell proliferation and differentiation. Using non-contact high-resolution Brillouin spectro-microscopy we showed that the corneal outer edge (limbus) has significantly lower bulk modulus compared to the central cornea, and that this difference is precisely delimited in the organ. Furthermore, the areas of the limbus with distinctly softer properties were shown to be associated with limbal epithelial stem cell (LESC) residence. Based on these findings, we then provided the first demonstration of the capacity to modulate LESC phenotype, both *in vivo* and *ex vivo*, solely through the recreation/restoration of suitable biomechanical niches. These results thus confirm the fundamental role of corneal biomechanics in directing epithelial stem cell behavior.

## Introduction

The function of the human cornea is largely dependent on the maintenance of a healthy stratified epithelium, which in turn relies upon a population of stem cells located in its periphery, the limbus (Lehrer et al., 1998). These limbal epithelial stem cells (LESCs) proliferate and differentiate to repopulate the central corneal epithelium, where cells constantly undergo maturation, stratification, and ultimately, shedding from the ocular surface. These events have been shown to be modulated by biochemical and biophysical factors (Di Girolamo, 2015; Yoon et al., 2014). However, the mechanisms underpinning the homeostatic process of LESC self-renewal and differentiation remain largely unclear (West et al., 2015). More recently, a number of studies have shown that the behavior of LESCs, like other stem cell types (Engler et al., 2006), is strongly influenced by their immediate mechanical environment. This notion is supported by the cellular stiffness of LESCs (Bongiorno et al., 2016), as well as by the distinct structure (Boote et al., 2005), composition (Mei et al., 2012), and compliance (Hjortdal, 1996; Last et al., 2012) of the extracellular matrix (ECM) across the cornea. In particular, the impact of substrate stiffness on corneal epithelial cell attachment and viability (Chen et al., 2012), proliferation (Jones et al., 2012), and mechanosensing (Foster et al., 2014) has been explored *in vitro*, using biomimetic surfaces with defined elastic moduli. These studies showed that corneal epithelial cells grown on relatively soft substrates are able to retain specific LESC markers whereas cells cultured on corresponding stiff substrates are disposed to differentiate (Foster et al., 2014; Jones et al., 2012). This body of work provides substantial evidence that, at least *in vitro*, substrate rigidity regulates LESC phenotype.

To understand if such regulation occurs *in vivo*, we presently built a high-resolution Brillouin spectro-microscope (Figure 1a) and used it to characterize the mechanical properties of live human corneas with unprecedented detail. This technique is based on the interaction of light with spontaneous acoustic phonons in the GHz frequency range. By measuring the optical frequency shift of the scattered light, Brillouin scanning can probe the local spontaneous pressure waves within biological tissues, from which the longitudinal modulus can be evaluated (Antonacci et al., 2013). Relating the hypersonic (GHz) modulus directly to the quasi-static modulus and Young’s modulus is complicated by acoustic dispersion that can increase moduli significantly at GHz frequencies. However, the technique does provide information about variations and changes in mechanical properties at submicron scale (Elsayad et al., 2016). Previously, Brillouin spectro-microscopy (BSM) has been used to evaluate mechanical properties of cells and tissues both *in vivo* (Scarcelli et al., 2015a) and *in vitro* (Antonacci and Braakman, 2016; Scarcelli et al., 2015b), particularly in the cornea (Lepert et al., 2016; Scarcelli et al., 2012). Our newly-developed instrument was designed with a single-stage virtually imaged phased array (VIPA) spectrometer in combination with an original wavefront division adaptive interferometric, and a piezoelectric actuator (Lepert et al., 2016) to give high extinction of the elastically-scattered light and allowing faster (0.01-1s) Brillouin scattering measurements. As such, we were able to obtain organ-wide scans of the human cornea, with both unprecedented detail and speed. Crucially, this achievement allowed us to identify critical biomechanical differences between the (softer) limbus and the (stiffer) central cornea, which were then correlated to the phenotype of corneal epithelial cells. This data supported our hypothesis that epithelial cell differentiation across the corneal surface is controlled by changes in substrate stiffness. More importantly, it allowed us to develop new a method to control the phenotype of corneal epithelial cells solely via modulation of the mechanical properties of collagen-based substrate materials.

**Figure 1:**
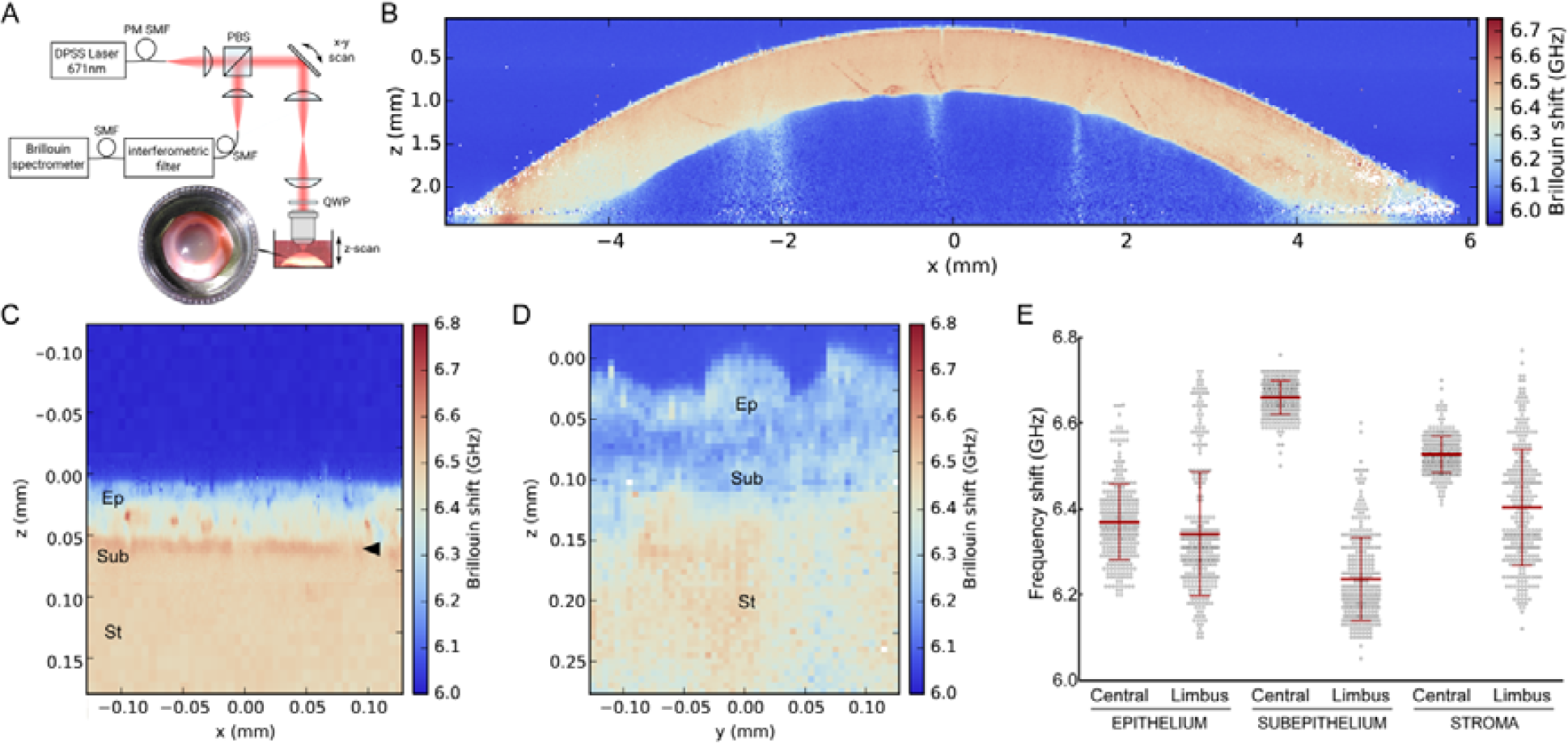
The corneal limbus has distinct mechanic characteristics compared to its center, as shown by the detailed, organ-wide characterization of the human cornea performed by Brillouin spectro-microscopy (BSM). **a**) Schematic representation of the Brillouin spectro-microscope (DPSS Laser, diode-pumped solid-state laser; PBS, polarizing beam-splitter; QWP, quarter-wave plate; PM-SMF, polarization-maintaining single-mode fiber), showing the confocal microscope, elastic scattering filter, VIPA spectrometer, and the sample in its immersion medium. Fresh human corneas were maintained intact after enucleation and kept immersed in dextran-containing Carry-C to preserve the cornea’s natural thickness, hydration, and transparency state (inset) during BSM measurements. **b**) Representative whole-organ *X*-*Z* scans of Brillouin frequency shifts from a healthy intact human cornea. Brillouin spectra were acquired with a sample spacing of 20 µm, over a 12 × 3 mm (*X-Z*) transverse section corresponding to the full corneal width and depth, respectively. **c**) Representative *Y*-*Z*-scan of Brillouin frequency shifts of central cornea performed every 2.5 µm, showing a distinct epithelium (*Ep*; depth = 0-50 µm), sub-epithelial layer (*Sub*; 50-65 µm), and stroma (*St*; >65 µm), as well as the location of the Bowman’s layer (*arrowhead*). **d**) Representative *Y*-*Z*-scan of Brillouin frequency shifts of corneal limbus performed every 5 µm, showing the epithelium (depth = 0-50/60 µm), sub-epithelial layer (60-120 µm), and stroma (>120 µm). **e**) Brillouin shift measurements performed through the epithelium, the sub-epithelial layer, and the stroma of both central cornea and limbus were evaluated quantitatively and compared statistically from three independent experiments (100 individual measurements/tissue/experiment), as well as the average ± S.D. (*n*=3; ** and *** corresponded to *p*<0.01 and 0.001, respectively). Shift value distribution was consistently similar between individual corneas but distinct in central *vs* limbus.

## Results and Discussion

### BSM shows clear transition between softer limbus and stiffer central anterior cornea

The new technological development of the BSM allowed whole-organ imaging of immersed human corneas (Figure 1b; Supplementary Figure 1a) to be performed through the entire 12 mm-span and 3 mm-depth of the cornea at high resolution (Figure 1b; 600 × 150 = 9×10^4^points) and accuracy (Supplementary Figure 1b). This approach facilitated the identification of numerous biomechanical features. First, it showed that the limbus had, overall, lower Brillouin frequency shift values, indicating it was more compliant than the central cornea (Figure 1b). Secondly, at very high resolution (2.5-5 µm distance between individual measurements), it identified the highest frequency shifts (i.e., stiffest tissue) in a discontinuous, 10-15 µm thick sub-epithelial layer of the central cornea (Figure 1c), probably corresponding to the Bowman’s layer (Last et al., 2012). Finally, it showed that this layer of stiff matrix was absent in the limbus (Figure 1d). The remarkable differences in Brillouin frequency shift between the anterior region of the central cornea and the limbus prompted a quantitative evaluation of these regions (Figure 1e). Brillouin transversal section scans in the central cornea showed a multi-layered epithelium approximately 50 µm thick (Figure 1c) with average ± S.D. frequency shift of 6.37±0.09 GHz (Figure 1e) and basal columnar cells presenting stiff nuclei, as previously predicted (Foster et al., 2014), followed by a 10-15 µm-thick Bowman’s layer and the anterior-most stroma with 6.66±0.04 and 6.53±0.04 GHz shifts, respectively (Figure 1e). In contrast, the limbal epithelium ranging 40-60 µm depth (Figure 1d) and 6.34±0.14 GHz was followed by matrix with significantly lower frequency shift (6.24±0.09 GHz), which in turn was followed by tissue exhibiting slightly higher shifts (6.40±0.14 GHz, respectively; Figure 1e), probably corresponding to the continuation of the corneal stroma under the limbus (Figure 1b) (Van Buskirk, 1989).

Consistently, the distinction in frequency shifts between epithelium and the sub-epithelial matrix immediately underneath was less clear in the limbus than in the central cornea (compare Figure 1c and d). The limbus also showed a broader distribution range of Brillouin frequency shift values in all its layers, whereas the central corneal was characterized by uniform biomechanical properties within each layer (Figure 1e), with a profile consistent to that from previous studies (Scarcelli et al., 2012). The clear distinction between the mechanically-heterogeneous limbus and the more regular central cornea was more evident in very high-resolution scans (Supplementary Figure 1c) and under alternative vantage points (Video 1 and 2). This distribution pattern was probably due to the furrowed topography of the limbus, and the presence of numerous structures such as the Palisades of Vogt, focal stromal projections, and limbal crypts and/or pits (Video 1), particularly abundant in the superior and inferior limbus but less so in the temporal and nasal side of the cornea (Di Girolamo, 2015; Lagali et al., 2013; Miri et al., 2012; Shortt et al., 2007) (Supplementary Figure 1d). Moreover, the very high-resolution Brillouin scans also identified multiple pocket regions within the limbal epithelium that exhibited significantly lower shifts compared to their immediate surrounds (Figure 1d; Supplementary Figure 1e). Three-dimension Brillouin scans showed that these pockets comprise multiple spherical units 10-12 µm in diameter (Video 2) that are compatible, dimension (Romano et al., 2003) and biomechanical-wise (Bongiorno et al., 2016), with LESCs surrounded by stiffer cells and matrix.

### BSM and immunofluorescence analysis shows LESCs residing on soft limbal matrix

Importantly, these limbal regions identified by BSM also represented the location of cells expressing ABCG2, CK-15, nuclear β-catenin, and ΔNp63 (Figure 2a-c; Supplementary Figure 1f), markers consensually associated with LESCs (Davies et al., 2009; Di Iorio et al., 2005; Nakatsu et al., 2011; Watanabe et al., 2004). In contrast, the CK3/12-positive epithelial cells from the central cornea were negative for LESC markers (Figure 2a-c), indicating that these constituted a more differentiated epithelium. The co-location of areas of low-Brillouin shift with the immuno-anatomical limbus was further confirmed by the geometry of its features (Supplementary Figure 1e-f), as well as by the expression of extracellular matrix components and corresponding cell receptors (Figure 2). The distribution of the low-Brillouin shift limbal sub-epithelial matrix (Figure 1e), classically identified as conjunctival stroma and Tenon’s capsule (Van Buskirk, 1989), was well correlated to the collagen-Ipositive/collagen-V-negative connective tissue immediately underlying the resident LESCs (Figure 2a-c). Moreover, the focal distribution of laminin-γ3, a characteristic marker of limbal basement membrane (Schlotzer-Schrehardt et al., 2007; Torricelli et al., 2013) along with the strong expression of integrin-α9, a transient amplifying cell/LESC marker (Schlotzer-Schrehardt and Kruse, 2005; Stepp, 2006), showed that this region corresponded to the corneal limbus (Figure 2d). The low-Brillouin shift values were also well correlated with the tissue’s structural features, previously characterized as comprising less compact collagen lamellae with irregularly arranged, branched, and intertwined collagen bundles (Boote et al., 2003; Komai and Ushiki, 1991). In contrast, the location of CK3-positive epithelial cells (Figure 2e) was well correlated to the regions of the central cornea with the highest-Brillouin shift values (Figure 1b). These regions showed to be comprised by ubiquitous corneal and conjunctival basement membrane components such as collagen-VII (Mei et al., 2012; Resch et al., 2009) and laminin-1 (Schlotzer-Schrehardt et al., 2007), but not laminin-γ3 (Figure 2de). Furthermore, the matrix under the central corneal epithelium was strongly positive for both collagen-I and collagen-V (Figure 2). This distinctive composition plays an important role on collagen fibril diameter and lamellar organization of the corneal stoma (Birk, 2001), which in turn has a critical influence in tissue transparency and elastic modulus (Boote et al., 2005; Hjortdal, 1996). Signal quantification also supported the distinctive expression pattern of markers in both limbus and central cornea (Figure 2f). Taken together, these results showed that, in the human cornea, LESCs populate tissues that are significantly softer compared to those supporting differentiated epithelial cells, thus constituting a niche with distinct biomechanical, as well as a biochemical/biomolecular profiles.

**Figure 2:**
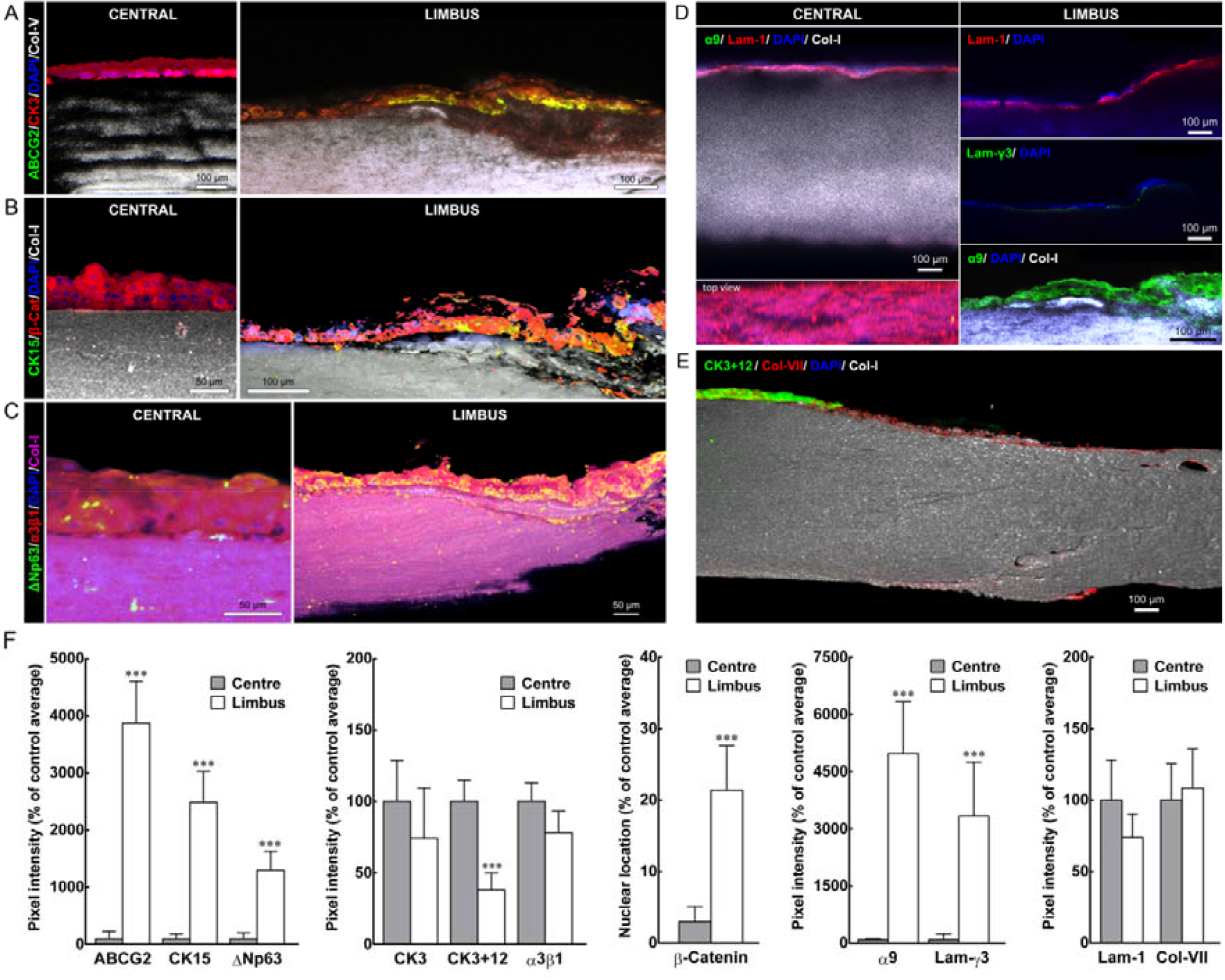
The location of the molecular markers of corneal epithelial stem cells corresponds to regions in the limbus with distinctly softer mechanical properties. Representative confocal immunofluorescence micrographs of corneal epithelial cell markers and ECM components were used to reconstruct in 3D the limbus and central cornea. Expression of LESC markers ABCG2 (**a**), CK15 (**b**), and ΔNp63 (**c**), were expressed by limbal epithelial cells supported by collagen-I-positive/collagen-V-negative matrix, but not by central corneal epithelium (**a**-**c**). Markers such as CK3 (**a**), β-catenin (**b**), and integrin-α3β1 (**c**) showed higher expression in the central corneal epithelium. Histochemical distinction between central cornea and limbus was further evidenced by the basement membrane markers and corresponding receptors. The limbus showed a discontinuous distribution of laminin-1 compared to the central cornea, and the specific expression of laminin-γ3 and integrin-α9 (**d**). Conversely, the central corneal epithelium was positive for CK3+12, as well as for collagen-VII (**e**). Cell nuclei were detected using DAPI. **f**) Marker expression was quantified and represented as average ± S.D. from three independent experiments (*n* = 3; *** corresponds to *p*< 0.001).

### Substrate stiffness controls corneal epithelial cell phenotype *in vitro*

The surprisingly distinct properties of the limbus observed by BSM reinforced our hypothesis that biomechanics plays a role in controlling the phenotype of LESCs (Foster et al., 2014). Conversely, the biomechanical features of the central cornea sustained the notion of a mechanical differential (i.e., gradient or step-change in substrate stiffness) driving corneal epithelial cell differentiation (Foster et al., 2014; Shin et al., 2013). It is then reasonable to assume that the measured differences in substrate stiffness across the cornea play an important role in the tissue’s function/homeostasis, by regulating the phenotype of corneal epithelial cells via mechanotransduction (i.e., maintaining undifferentiated cells in the limbus, and promoting their differentiation in the central cornea).

In this perspective, we aimed at further investigating the response of LESCs to surface compliance, first using an *in vitro* model. Previously, high-density collagen-I gels (tissue mimics) with different levels of stiffness via plastic compression have been used to that purpose (Gouveia et al., 2014). Now we explored the premise that collagen gel stiffness could be affected through its partial digestion using a type-I collagenase. This approach was chosen as it would also facilitate the topical application on corneal tissues *in vivo*, using a formulation currently approved for connective tissue softening (Brunengraber et al., 2014; Gelbard et al., 2013; Hurst et al., 2009). Thus, we used plastic-compressed collagen gel discs 2.5 cm in Ø, to which a solution of collagenase was applied in discrete areas for up to 60 min at 37°C (Figure 3a). Treated gels were shown to be partially digested, with enzymatic cleavage restricted to the collagenase-soaked areas (Figure 3b, treated *vs* untreated), and acting through the entire thickness of the matrix without reducing gel thickness or compromising its structural integrity (Figure 3b).

**Figure 3:**
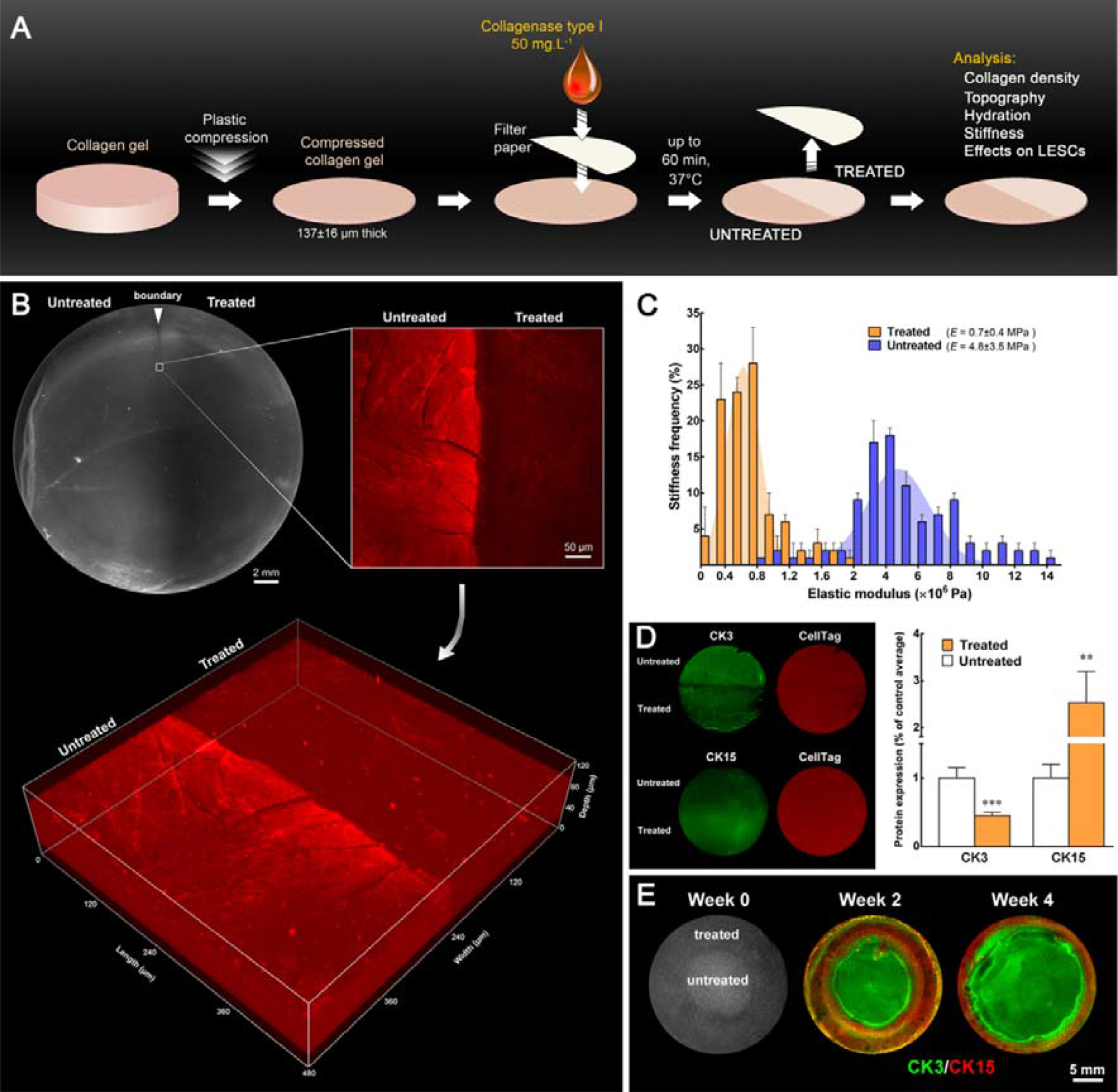
LESC phenotype can be controlled through fine modulation of the mechanical properties of collagen substrates. **a**) Schematic representation of the collagenase treatment method used to modulate the stiffness of compressed collagen gels. High-density, plastic-compressed collagen gels were softened with collagenase type-I solution in well-defined areas (ring-shaped, semi-circular, or entire gel surface) for up to 60 min. **b**) Analysis of compressed collagen gel density after collagenase treatment. The regions corresponding to collagenase-treated collagen gels showed increased transparency under bright-field imaging (upper left panel; scale bar, 2 mm) and lower collagen density compared to untreated regions, as indicated by the lower collagen-I detection by immunofluorescence confocal microscopy (upper right panel; scale bar, 50 µm). The confocal *Z*-scans (lower panel) demonstrated that the difference in signal intensity between treated and untreated areas was not restricted to the surface, indicating that the collagenase in solution acted through the entire depth of the compressed collagen maintaining a defined treatment zone. **c**) Average frequency ± S.D. of the elastic modulus, *E* (MPa), of treated (orange) and untreated gels (blue bars) calculated from three independent experiments using force-distance spectroscopy (*n*=3). The frequency histograms of treated and untreated gels were used to calculate Gaussian curves by non-linear regression (orange and blue areas, respectively), with corresponding *E* = 0.7±0.4 and 4.8±3.5 MPa. **d**) Effects of substrate stiffness on the expression of CK3 (differentiation) and CK15 (LESC protein marker) in hLECs grown for 4 weeks on treated and untreated regions of collagen gels (green staining) after normalization for total cell number (red staining), and represented as average ± S.D. from three independent experiments (*n* = 3; ** and *** corresponds to *p*<0.01 and 0.001, respectively). **e**) Creation of a pseudo-limbus. Cells growing on ring-shaped softened areas (week 0) expressed higher levels of CK15 compared to the high CK3-positive cells growing on the untreated (stiffer) central region of the collagen gels, up to 4 weeks in culture.

The effect of collagenase treatment on collagen gel topography, hydration, and stiffness was further characterized (Supplementary Figure 2a-e). Particularly, gels treated for 60 min had *E* = 0.7±0.1 MPa, almost an order of magnitude lower compared to that of mock-treated gels, with *E* = 5.2±1.2 MPa, corresponding to a statistically significant difference (*p* = 0.002) in frequency distribution of *E* values (Figure 3c). The different compliance between treated and untreated gels constituted an important result, as it was subsequently shown to affect corneal epithelial cell phenotype. Human limbal epithelial cells (hLECs) previously isolated from human corneal limbal ring explants were grown on softened (treated) collagen gels, where they showed significantly higher proliferation (*p* = 0.048) (Supplementary Figure 2f) and slower migration speed (*p* = 0.0003) compared to cells grown on the stiffer, untreated gels (Supplementary Figure 2g). In addition, hLECs grown on softened gels showed a significant decrease in CK3 (*p* = 0.0004) and increase in CK15 expression (*p* = 0.0012) at both transcriptional (Supplementary Figure 3a) and protein levels (Figure 3d), in a particularly well-defined pattern delimited by the boundaries of enzyme pre-treatment (Figure 3d; left panel). Furthermore, cells grown on collagenase-treated gels showed a lower, albeit not significantly different expression of inflammation-related markers (Supplementary Figure 3b). These results indicated that the enzymatic digestion can produce precisely controlled, localized differences in the stiffness of high-density collagen gels, resulting in the removal of up to one third of the original collagen content from within the gels without altering their volume or micro-topography, and subsequently maintain hLECs undifferentiated. As such, we then used the collagenase treatment to create a ring of softer collagen (representing the limbus, having shown by BSM that this tissue is naturally significantly softer) surrounding a stiffer area (representing the central cornea). Cells subsequently grown on the treated, outer-ring area (pseudo-limbus) expressed higher levels of CK15 up to 4 weeks, whereas hLECs on the central, untreated region of the gels displayed higher CK3 expression (Figure 3e). Furthermore, the maintenance of CK15 expression by hLECs on collagen gels was, overall, significantly higher than on tissue culture plastic at equivalent stages in culture (Supplementary Figure 3c). These results suggested that cells residing upon this collagenase-created pseudo-limbus retained a more undifferentiated, LESC-type phenotype. Together, these results demonstrate that the application of collagenase to high-density collagen gels creates localized areas that act as a niche for hLECs to maintain LESC properties such as those observed in the corneal limbus. The notion that LESC phenotype maintenance depended on the mechanical properties of softened gels and not on interactions with new cues created after collagenase treatment was supported by previous work using hLECs on semi-compressed collagen gels (Foster et al., 2014). The topographical, mechanical, and phenotype-modulating properties of semi-compressed collagen gels, despite being comparable to those of enzyme-softened gels (Supplementary Figure 3d-g), were solely controlled by the time and load of plastic compression (Jones et al., 2012), and therefore independent of exposed cryptic epitopes. Collagenase presented also the advantage to allow softening of collagen-based tissues both *in vivo* and *ex vivo*, and could thus be applied to evaluate the ‘cell phenotype-through-biomechanical modulation’ strategy demonstrated above, but using live, natural tissues (e.g., the cornea).

### Softening the human central cornea creates a limbus-like mechanical and phenotypic milieu

In order to investigate the ability to control cell phenotype via modulation of the biomechanical properties of natural tissues, collagenase was used to soften limited and well-defined areas in the center of freshly enucleated human corneas (Supplementary Figure 4a). This experiment aimed at testing if alterations in the mechanical properties of the central cornea towards a more compliant, limbus-like environment, would allow creating an alternative mechanical niche suitable for LESCs. As predicted from the plastically-compressed collagen gel study, topical collagenase application to the human cornea resulted in lower Brillouin frequency shifts in the collagenase-treated, but not in mock-treated areas of the cornea (Figure 4a), with no loss of accuracy (Supplementary Figure 4b). Specifically, the collagenase treatment significantly lowered the Brillouin frequency shift in the central sub-epithelial and anterior stroma matrix compared to controls (Figure 4b) while surprisingly maintaining the epithelium layers in place, albeit in a less evenly distributed pattern (Figure 4b, right panel). Notably, the Brillouin frequency shift from the treated sub-epithelial matrix of the central cornea (6.45±0.13 GHz) was statistically similar to that of the sub-epithelium from the control (untreated) limbus (6.34±0.07 GHz) (Figure 4c). This indicated that collagenase treatments can soften the central cornea towards a limbus-like level of compliance. Collagenase further reduced the stiffness of the limbus (Supplementary Figure 4c), namely at the sub-epithelium level (Figure 4c).

**Figure 4:**
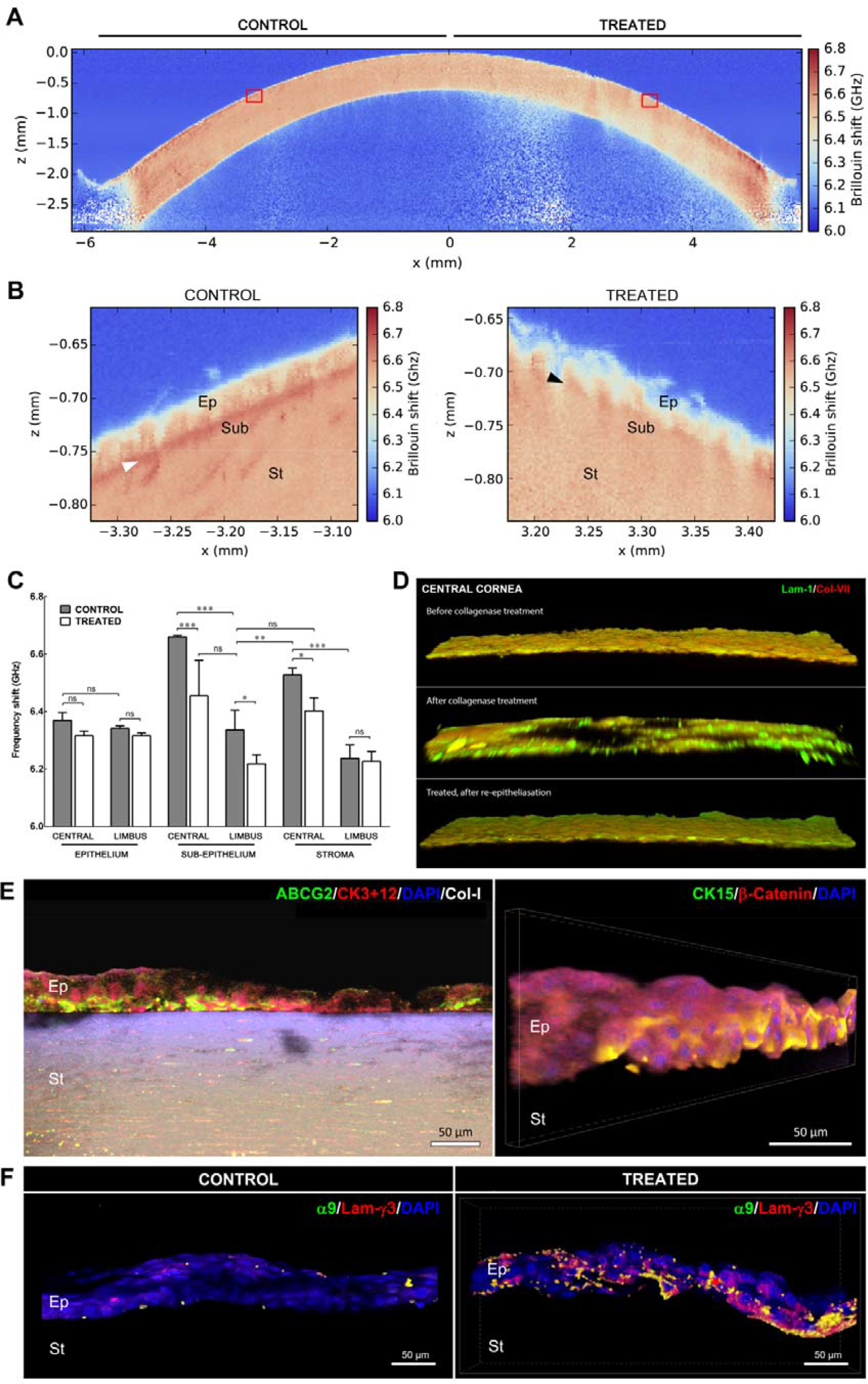
Softening of corneal tissue with collagenase increases expression of LESC markers during *ex vivo* re-epithelialization. **a**) Representative whole-cornea *X*-*Z* scan of Brillouin frequency shifts measured at a sample spacing of 20 µm from healthy intact human corneas after collagenase treatment. Insets correspond to (**b**) the regions of the central cornea softened with collagenase (treated) or left untreated (control) analyzed at very high-resolution scanning, using a sampling distance of 2.5 µm, showing a distinct epithelium (*Ep*), sub-epithelial layer (*Sub*), and stroma (*St*), as well as the location of the Bowman’s layer (*white arrowhead*). This detailed analysis evidenced the loss of the stiffer Bowman’s layer (*black arrowhead*) and reduced Brillouin frequency shifts in both epithelium and stroma resulting from collagenase treatment. **c**) Quantification of Brillouin frequency shifts from tissues of the anterior part of both treated (*white*) and control regions (*grey bars*) of the central cornea and limbus. The plot represents average ± S.D. of measurements taken from the epithelium, sub-epithelium, and stroma from three independent experiments (100 individual measurements/tissue/experiment; *n*=3; *, **, and *** represented *p*<0.05, 0.01, and 0.001, respectively). **d**) Representative confocal immunofluorescence micrographs (3D reconstruction) of laminin-1 (green) and collagen-VII distribution (red staining) in the central cornea, before (upper) and after collagenase treatment (central panel), and after reepithelialization (lower panel). **e**) Representative confocal immunofluorescence micrographs (3D reconstruction) of collagenase-softened central cornea after re-epithelialization. Epithelial cells repopulating softened corneas expressed ABCG2 and CK15 (LESC markers; green) while showing lower CK3+12 and β-catenin expression (differentiation markers; red staining) compared to cells growing on the stiffer, untreated corneas (control). **f**) Epithelial cells expressed integrin-α9 and deposited laminin-γ3 when grown on collagenase-softened corneal substrates, but not on untreated central cornea (control). Cell nuclei were detected using DAPI. Scale bars, 50 µm.

The altered mechanical properties of the collagenase-treated central cornea were in part derived from the degradation of the collagen-rich matrix of the anterior stroma, including the Bowman’s layer (Figure 4b). However, confocal immunofluorescence analysis showed that the basement membrane in the central cornea was also affected, presenting large gaps in the distribution of collagen-VII and laminin-1 following collagenase treatment (Figure 4d). Nonetheless, after careful debridement through washing, both treated and control central corneas successfully served as *ex vivo* growth substrate for hLECs, supporting reepithelialization (Figure 4e) and basement membrane re-deposition (Fig. 4d). Similarly to their behavior on collagen gels, cells grown on softened areas of the central cornea continued to express LESC markers such as ABCG2 and CK15, particularly at the basal layers of the new epithelium (Figure 4e). Moreover, these cells deposited focalized laminin-γ3 while also expressing integrin-α9 (Figure 4f), both characteristic markers of the limbal niche (Schlotzer-Schrehardt et al., 2007; Schlotzer-Schrehardt and Kruse, 2005; Stepp, 2006; Torricelli et al., 2013). In contrast, hLECs grown on mock-treated areas of the central cornea failed to express any of these markers. Signal quantification showed that this differential expression was significant for all markers tested (Supplementary Figure 4d). Taken together, these results showed that collagenase treatment has the potential to soften the matrix of the anterior layers of the central cornea, and thus create a substrate that is mechanically similar to the natural limbus, and biologically capable of maintaining a LESC-like phenotype.

### Softening the rabbit central cornea modulates epithelial cell phenotype

Next, we used a rabbit corneal model to demonstrate that phenotype-through-biomechanical modulation of corneal epithelial cells can be controlled *in vivo* through precise application of collagenase (Figure 5). The outcome of collagenase treatment on intact and debrided (i.e., epithelium removed) rabbit corneas was followed by clinical observation, slit-lamp biomicroscopy, and immunohistochemistry analyses 1 and 5 days post-intervention (Figure 5a; Supplementary Figure 5). The *in vivo* softening of intact corneas after collagenase treatment elicited a radical change in epithelial cell phenotype, with molecular markers showing major differences in expression compared to corresponding controls (mock-treated rabbit corneas; Figure 5b), in line with the effects obtained *in vitro* (with collagen gels) and *ex vivo* (with organ-cultured human cornea). Specifically, epithelial cells residing on the softened central cornea were shown to express CK15, ABCG2, ΔNp63, laminin-γ3, and integrin-α9 five days after treatment (Figure 5b; Supplementary Figure 5 and 6). Conversely, softening the central cornea induced a reduction in CK3/12 expression in the epithelium and resulted in a partially-degraded basement membrane, as shown by the reduced levels of collagen-IV and collagen-VII (Figure 5b; Supplementary Figure 5 and 6). These differences in expression were significant (Figure 5b), and represented important alterations in the normal phenotype of epithelial cells from the central cornea. The softening of intact corneas also maintained corneal integrity and transparency (Figure 5a), as well as preventing any substantial irritation, inflammation, edema, or change in intraocular pressure (Supplementary Table S1). Moreover, no conspicuous neo-vascularization was observed following collagenase treatment of the cornea either on a macroscopic (Figure 5a) or microscopic level (Supplementary Figure 7). The maintenance of the epithelial barrier function, even after collagenase treatment (Supplementary Figure 8), was probably the main factor protecting intact corneas from these deleterious effects (Figure 5a).

**Figure 5:**
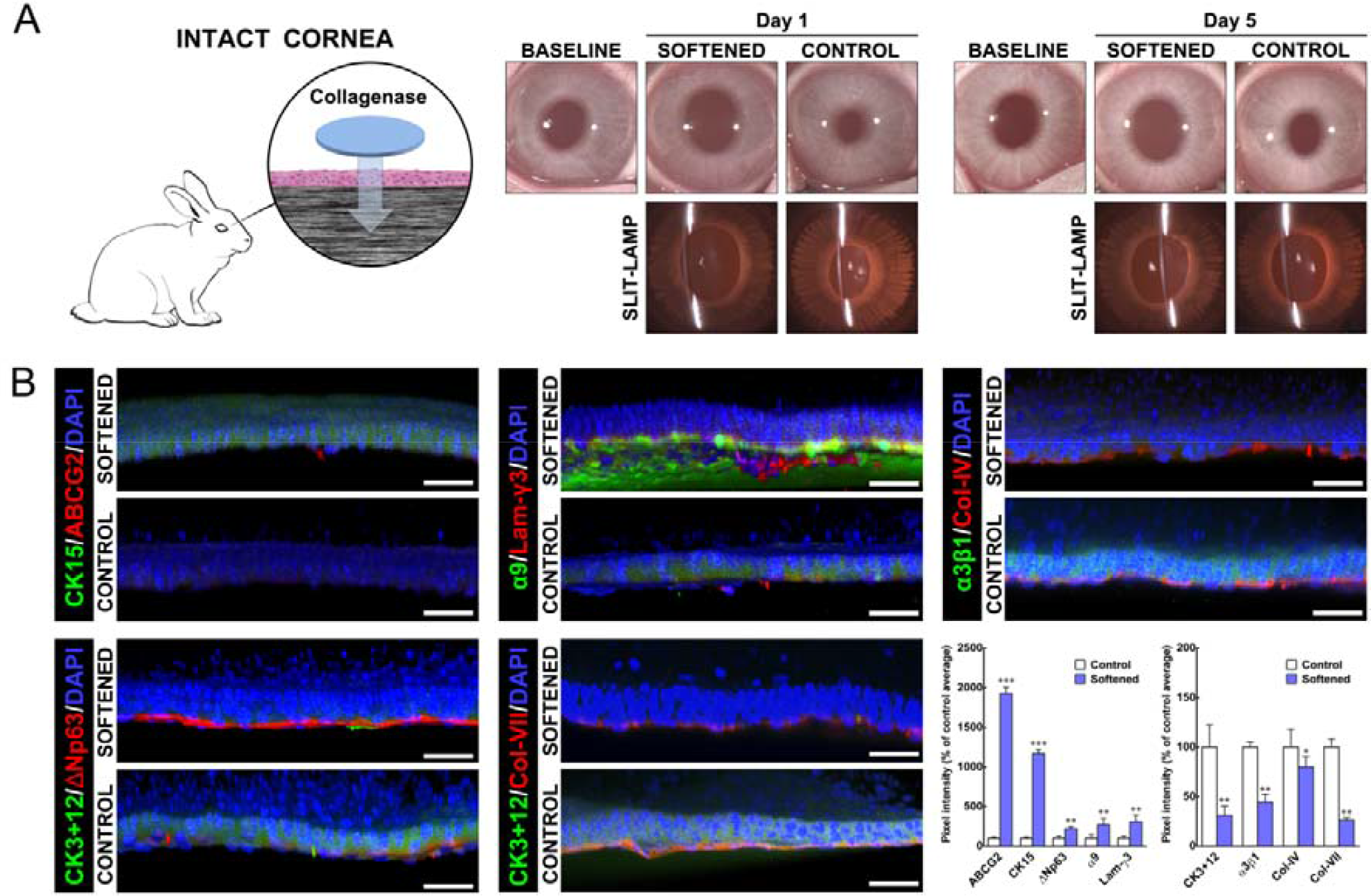
Softening of central corneal tissue with collagenase increases expression of LESC markers *in vivo*. **a**) Schematic representation of the collagenase treatment method used to soften the central region of intact corneas in live rabbits. Clinical observation and slit-lamp examination was performed 1 and 5 days post-intervention, and compared to pre-intervention results (baseline). **b**) Representative confocal immunofluorescence micrographs (3D reconstruction) of central corneal epithelium 5 days after collagenase treatment (softened) and corresponding quantification of marker expression. Cells on softened corneas expressed higher levels of ABCG2, CK15, ΔNp63, integrin-α9, and laminin-γ3 (LESC-characteristic markers) and lower levels of CK3+12 and integrin-α3β1 (differentiation markers) compared to cells growing on the stiffer, untreated corneas (control). Collagen-IV and collagen-VII were also detected in collagenase-treated (softened) corneas, albeit at lower levels compared to control. Cell nuclei were detected using DAPI. Scale bars, 50 µm. Marker expression was represented as average ± S.D. from three independent experiments (*n* = 3; *, **, and *** corresponds to p<0.05, 0.01, and 0.001, respectively).

The fact that collagenase-treated tissues were able to retain their shape and structure without eliciting a major pro-inflammatory response (Supplementary Figure 3b) is of great importance for the validity of this method. Specifically, it showed that the collagen matrix can be softened without compromising the integrity of the cornea. Moreover, it supported the notion that the modulation in corneal epithelial cell phenotype was due to alterations in tissue biomechanics, and not the result of inflammation-related signaling. This is important, as severe inflammation of the ocular surface has been associated with corneal injury and disease leading to LESC loss (Deng et al., 2012), namely through mechanotransduction pathways (Nowell et al., 2016). These results thus indicated that the collagenase treatment was innocuous, if not beneficial, to the culture of corneal epithelial cells on collagen-rich matrices.

### Collagenase treatment restores limbus capacity to support LESCs after stiffening

Finally, we explored the effect of collagenase on the phenotype of corneal epithelial stem cells grown on artificially stiffened corneas. An alkali burn technique was thus developed, as this method allowed the experiments to be performed in small, well-defined, localized areas. First, human corneas subjected to alkali burn *ex vivo* were treated with collagenase and tested for their mechanical properties, as well as their ability to support the LESC phenotype (Figure 6). Not unexpectedly, the alkali burn led to major alterations in the corneal limbus, with clearance of most of its non-fibrillar elements and significant stiffening of its matrix (Figure 6a and b). These effects were probably due to proteoglycan removal following alkali exposure (Scott and Thomlinson, 1998). Subsequently, alkali-burned corneas also failed to maintain the LESC phenotype of repopulating cells. Specifically, hLECs seeded on the limbus of alkali-burned corneas showed significantly reduced expression of LESC markers ABCG2, CK15, ΔNp63, and integrin-α9, while expressing higher levels of differentiation marker CK3 compared to cells on non-burned, control tissue (Figure 6c and d). However, these effects were shown to be reversed by application of collagenase on alkali-burned corneas, with treatment leading to reduced collagen fiber density and significant decrease in the stiffness of the limbus matrix (Figure 6a and b). But most importantly, the recovery of the original tissue compliance was accompanied by the ability to support re-epithelialization by hLECs expressing LESC markers (Figure 6c and d).

**Figure 6:**
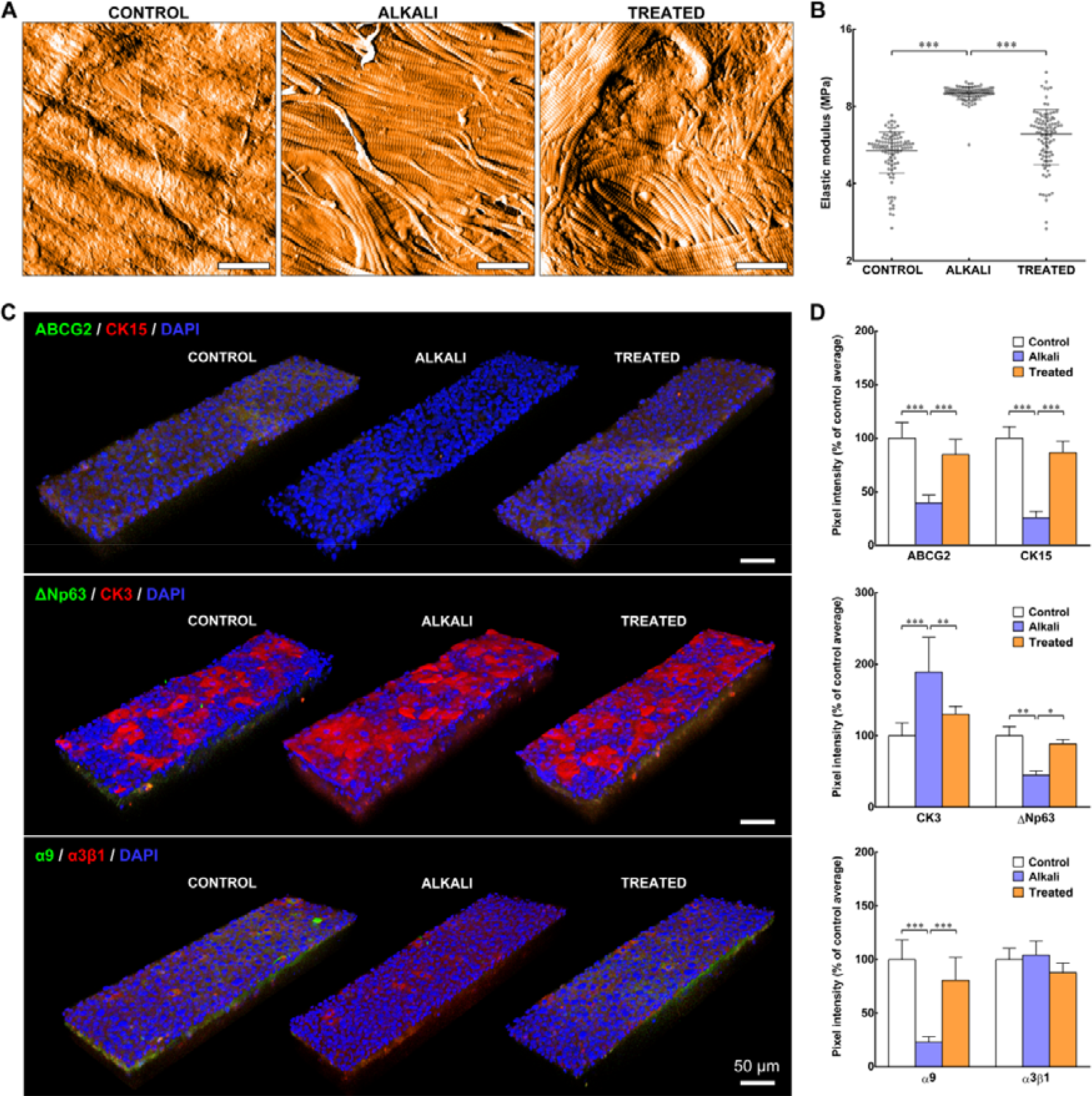
Softening of alkali-burned corneal tissue with collagenase restores expression of LESC markers *ex vivo*. **a**) Representative topography of limbal sub-epithelial matrix after application of PBS (control), 0.5 M NaOH (alkali), or NaOH followed by collagenase softening (treated) of whole human corneas, analyzed by atomic force microscopy (AFM). AFM scans showed clearance of ECM components other than collagen fibers in tissue subjected to alkali burn, and reduced collagen fiber density after collagenase treatment. Scale bars, 1 µm. **b**) Force-distance spectroscopy analysis showed that the burn (alkali) significantly stiffened the limbal sub-epithelial matrix, and that subsequent collagenase softening (treated) restored the mechanical properties of the original tissue (control). The graph represents the distribution of calculated values of elastic modulus, *E* (MPa), and corresponding averages ± S.D. from three independent experiments (*n* = 3; *** corresponds to *p*< 0.001). **c**) Representative confocal immunofluorescence micrographs (3D reconstruction) of re-epithelialised limbus after alkali burn and collagenase treatment, with (**d**) corresponding marker expression quantification. Epithelial cells repopulating the alkali-burned limbus tissue *ex vivo* expressed significantly lower levels of LESC markers ABCG2, ΔNp63, CK15, and integrin-α9 while showing higher expression of CK3 differentiation markers compared to cells growing on control tissue. However, collagenase treatment successfully restored the ability of the alkali-burned limbus to support cells as in control tissues. Cell nuclei were detected using DAPI. Marker expression was represented as average ± S.D. from three independent experiments (*n* = 3; *, **, and *** corresponds to p<0.05, 0.01, and 0.001, respectively). Scale bars, 50 µm.

Similar results were observed in an *in vivo* chemical burn model. Rabbit corneas subjected to a localized alkali burn had the temporal half of the limbus damaged while maintaining its nasal half intact, to allow re-epithelialization (day 0, Supplementary Figure 9). On the second day post-burn, the affected areas of the limbus were either treated with collagenase (treated) or remained untreated (alkali), and were subsequently compared to undamaged limbus tissue (control). Immediately after the burn, the affected area of the limbus became opaque (Figure 7a). In untreated corneas, this haze showed little or no reduction in size (Figure 7b, *alkali*). In contrast, collagenase-treated tissues showed a significant (*p* = 0.002) reduction of haze area compared with alkali-burn tissues (*p* = 0.0023) and with time (90±3 to 56±5% of initial size from day 2 to 7, respectively) (Figure 7b). Furthermore, treating the burned limbus with collagenase allowed the tissue’s native cellular profile to be restored (Figure 7c). Specifically, the new epithelium populating the collagenase-treated burned limbus tissue showed significantly higher expression of LESC markers CK15, ΔNp63, ABCG2, and integrin-α9, and lower expression of differentiation markers CK3 and α3β1 compared with their untreated (alkali) tissue counterparts, and an expression profile similar to the undamaged (control) limbus (Figure 7c and d).

**Figure 7:**
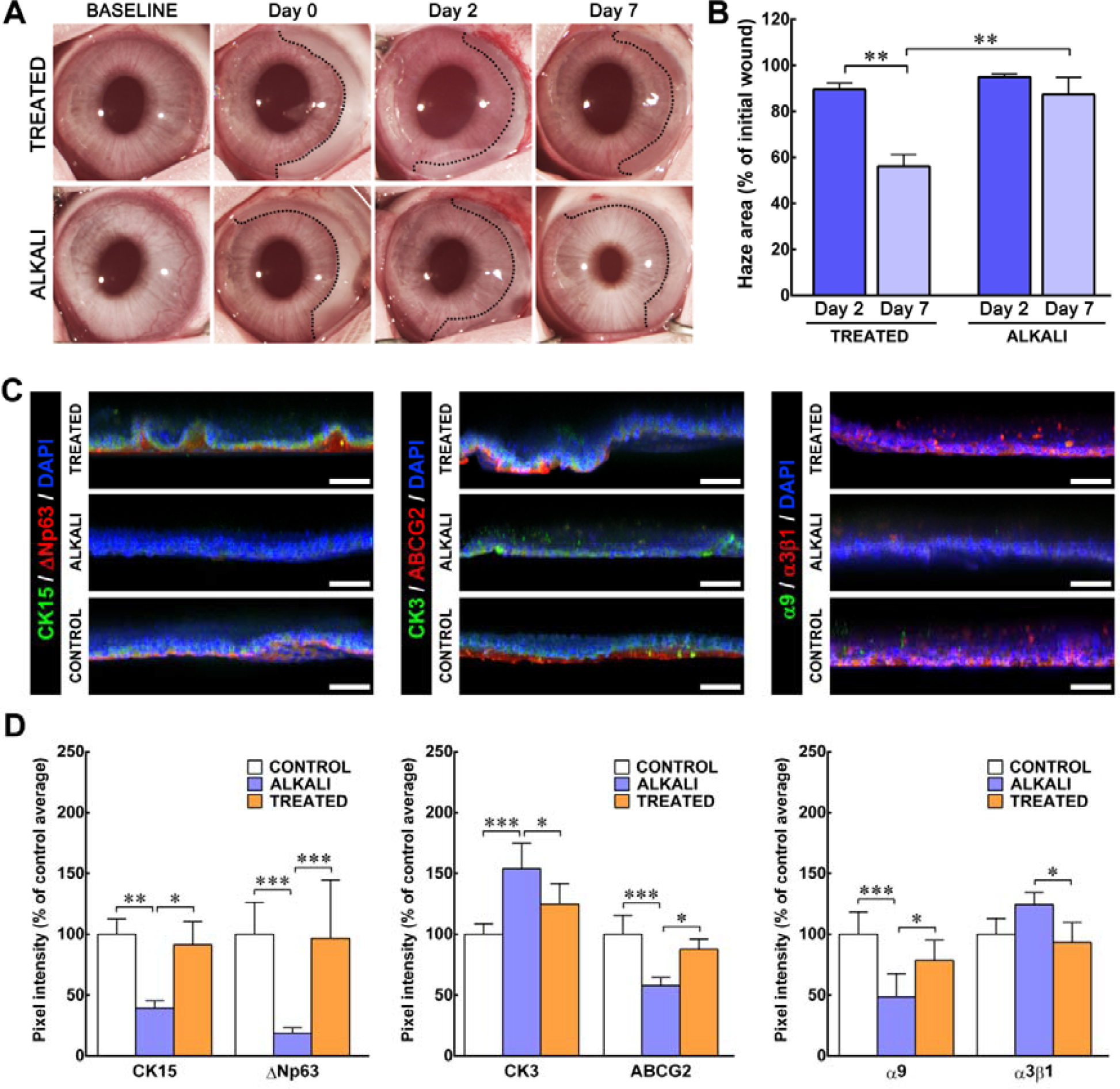
Treatment of alkali-burned limbus with collagenase restores expression of LESC markers *in vivo*. **a**) Representative images of chemically-burned rabbit corneas (alkali) and of alkali-burned corneas receiving collagenase treatment (treated), before (baseline) and after burn (day 0), as well as at day 2 and 7 post-burn. The haze resulting from the burn was delimited (traced lines), and its area quantified (**b**) at day 0 (initial wound), 2 (dark blue) and 7 (light blue bars). Values corresponded to haze area average ± S.D. from three independent experiments (*n* = 3; ** corresponds to *p* < 0.01. **c**) Representative confocal immunofluorescence micrographs (3D reconstruction) of chemically-burned (alkali) and collagenase-treated burned limbus (treated) at day 7, with (**d**) corresponding marker expression quantification. Epithelial cells repopulating the alkali-burned limbus *in vivo* expressed significantly lower levels of LESC markers CK15, ΔNp63, ABCG2, and integrin-α9 while showing higher expression of the CK3 differentiation marker compared to the undamaged limbus (control) tissue. However, collagenase treatment successfully restored the ability of the burned limbus to support cells as in control tissues. Cell nuclei were detected using DAPI. Marker expression was represented as average ± S.D. from three independent experiments (*n* = 3; *, **, and *** corresponds to *p*<0.05, 0.01, and 0.001, respectively). Scale bars, 50 µm.

Overall, these results showed that collagenase effectively reversed the stiffening of the corneal limbus following alkali burn and allowed the recovery of its natural LESC-supporting capacity. Moreover, the restorative effect of collagenase did not elicit substantial neovascularization, nor did it compromise the central cornea epithelium (Supplementary Figure 10). As such, this data demonstrated that the phenotype of corneal epithelial cells, and particularly of LESCs, can be influenced *in vivo* via the manipulation of their mechanical environment.

## Conclusions

In summary, this study demonstrates the strong correlation between compliance of the corneal tissue and epithelial cell phenotype. Specifically, the results show that compliant substrates support the growth of undifferentiated LESCs, whereas stiffer substrates promote their differentiation. Moreover, we applied this phenomenon to the development of a novel collagenase-based method to affect LESC phenotype-through-biomechanical modulation both *in vivo* and *ex vivo*. This work opens multiple areas of research, particularly regarding the use of BSM to study the mechanobiology of stem cell niches, in real time and with unparalleled detail and resolution, as well as the use of enzymes to regulate stem cell phenotype though modulation of the mechanical properties of substrate tissues *in situ*.

## Acknowledgments

This work was supported by the Medical Research Council (MRC-UK) research grant MR/K017217/2.

## Author Contributions

Conceptualization, R.M.G., C.P. and C.J.C.; Methodology, R.M.G., R.R.M., C.P. and C.J.C.; Investigation, R.M.G, G.L. and S.G.; Writing - original draft, R.M.G., C.P. and C.J.C.; Funding Acquisition and Supervision, R.R.M, C.P. and C.J.C.

## Declaration of Interests

The authors declare no competing interests.

## Experimental Procedures

### Brillouin spectro-microscopy

The design and setup of the Brillouin spectro-microscope and VIPA spectrometer, as well the evaluation method for the Brillouin scattering from fresh human corneas were performed as previously described (Lepert et al., 2016). Briefly, whole human corneas surrounded by a scleral ring were obtained from the National Disease Research Interchange (NDRI, RRID:SCR_000550) from cadaveric donors (ages ranging 42–74 years old with no prior history of corneal diseases or ocular trauma), in accordance with the Newcastle University and Newcastle-upon-Tyne Hospital Trust Research Ethics Committees’ guidelines. Isolated tissues were kept at room temperature in Carry-C preservative medium (Alchimia, Italy) during transport and Brillouin spectro-microscopy analysis to maintain the cornea’s natural hydration state. The measured Brillouin shifts were obtained by fitting the Stokes and anti-Stokes components of the spectra with Lorentzian function line profiles using non-linear least squares (SciPy optimized curve fit), and the amplitude of the fitted Lorentzian giving the strength of the spectral signal. The Brillouin shift for the backscatter geometry was given by 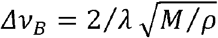 where *λ* is the optical wavelength in the sample, and and correspond to the longitudinal modulus and density of the sample, respectively (Lepert et al., 2016).

### Collagen gel production

Plastic-compressed high-density collagen gels were produced as previously described (Brown et al., 2005; Foster et al., 2014; Jones et al., 2012). Briefly, collagen gels were made by neutralizing sterile rat-tail type-I collagen (2.2 mg.ml^-1^ in 0.6 % acetic acid; First Link Ltd, UK) in 10× Modified Eagle’s Medium (MEM; Thermo Scientific, RRID:SCR_008452) and 1 M sodium hydroxide (Sigma-Aldrich, RRID:SCR_008988) at a 8:1:1 volume ratio. The solution was gently mixed and cast into circular molds (2 mL per 1.9 cm^2^ ø wells in 12-well plates) prior to gelling for 30 min at 37 °C. Collagen gels were then compressed between two layers of nylon mesh (50 µm mesh size) under a fixed load of 134 g for 5 min, or semi-compressed with 64 g for 2.5 min at room temperature. Compressed collagen gels were also coated with the matrix extracted from human corneas. Briefly, stromal tissues isolated from either the central cornea or limbus regions were minced and digested with 1 × 10^4^ activity units of collagenase per L^-1^ of PBS for 60 min at 37°C. Stroma and limbus protein extracts were then precipitated by incubation with 4× volume of ice-cold ethanol overnight at −20°C, followed by centrifugation at 10,000 ×*g* for 10 min at 4°C, and resuspended in PBS. The different corneal extracts were drop-spotted onto compressed collagen gels, and allowed to dry overnight to create a continuous coating.

### Collagenase treatments

Tissue softening was performed by applying collagenase type-I isolated from *Clostridium histolyticum* (#17018-029, Thermo Scientific). Dose and time of incubation were optimized to reproduce the biomechanical and functional properties of semi-compressed collagen gels^12^. Briefly, lyophilized collagenase powder was solubilized in phosphate buffered saline (PBS) at 5 × 10^-2^ g.L^-1^ (1 × 10^4^ activity units.L^-1^) and then applied onto sterilized Whatman Grade 1 filter paper until saturation, i.e., approximately 0.2 mL.cm^-2^ of paper. The collagenase-soaked paper cut-outs were then gently applied onto the surface of dense collagen gels and incubated for up to 60 min at 37°C. Treatment of whole human corneas was performed for 60 min at 37°C (2 units.cm^-2^.h^-1^ of total collagenase activity). The collagenase-soaked filter paper cut-outs were subsequently discarded, and the gels or corneas were washed thrice with PBS for 15 min on a rocker agitator to remove remaining collagenase enzyme and products from digestion. PBS-soaked filter paper cut-outs were incubated for 1 h using a similar method, in order to produce mock-treated specimens (treatment time = 0 min). Collagen gels treated for different durations were imaged by bright-field photography using a Nikon DSRL digital camera (Nikon, Japan) immediately after washes.

### Confocal immunofluorescence microscopy

Collagen gels were washed in PBS, blocked in PBS supplemented with 5% bovine serum albumin (BSA) and 0.1% Triton X-100 for 1 h, incubated overnight at 4°C with rabbit anti-collagen type-I primary antibody (Table 1) diluted 1:500 in blocking buffer, washed vigorously 4 × 15 min in PBS, and incubated in the dark for 2 h at room temperature with Alexa 594-conjugated goat anti-rabbit secondary antibody (Thermo Scientific) diluted 1:1000 in blocking buffer. After another 4 × 15 min washes with PBS, gels were immersed in Vectashield anti-fade medium (Vector Laboratories, UK), mounted on glass slides, and imaged by confocal fluorescence microscopy using a Nikon A1, with 1 µm-thick optical sections. Fresh human corneal tissues were fixed in 4% paraformaldehyde in PBS for 20 min and then sectioned in thick slices (0.5-0.8 mm) encompassing the central cornea, limbus, and sclera. Individual corneal slices were then blocked in PBS supplemented with 5% BSA and 0.1% Triton X-100 for 3 h, incubated overnight at room temperature with primary antibodies (Table 1) diluted 1:500 in blocking buffer, washed 4 × 1 h in PBS, incubated in the dark for 4 h at room temperature with the corresponding secondary antibodies and DAPI (Thermo Scientific) diluted 1:1000 in blocking buffer, washed again 4 × 1 h in PBS, and mounted onto glass slides. Data was analyzed using the NIS-Elements (RRID:SCR_014329) and the ImageJ v.1.7 software suite (RRID:SCR_003070), with quantification of expression performed by evaluating pixel intensity for each independent channel, and normalized against percentage of average pixel intensity of corresponding control. Experiments were performed three independent times (*n* = 3).

**Table 1:**
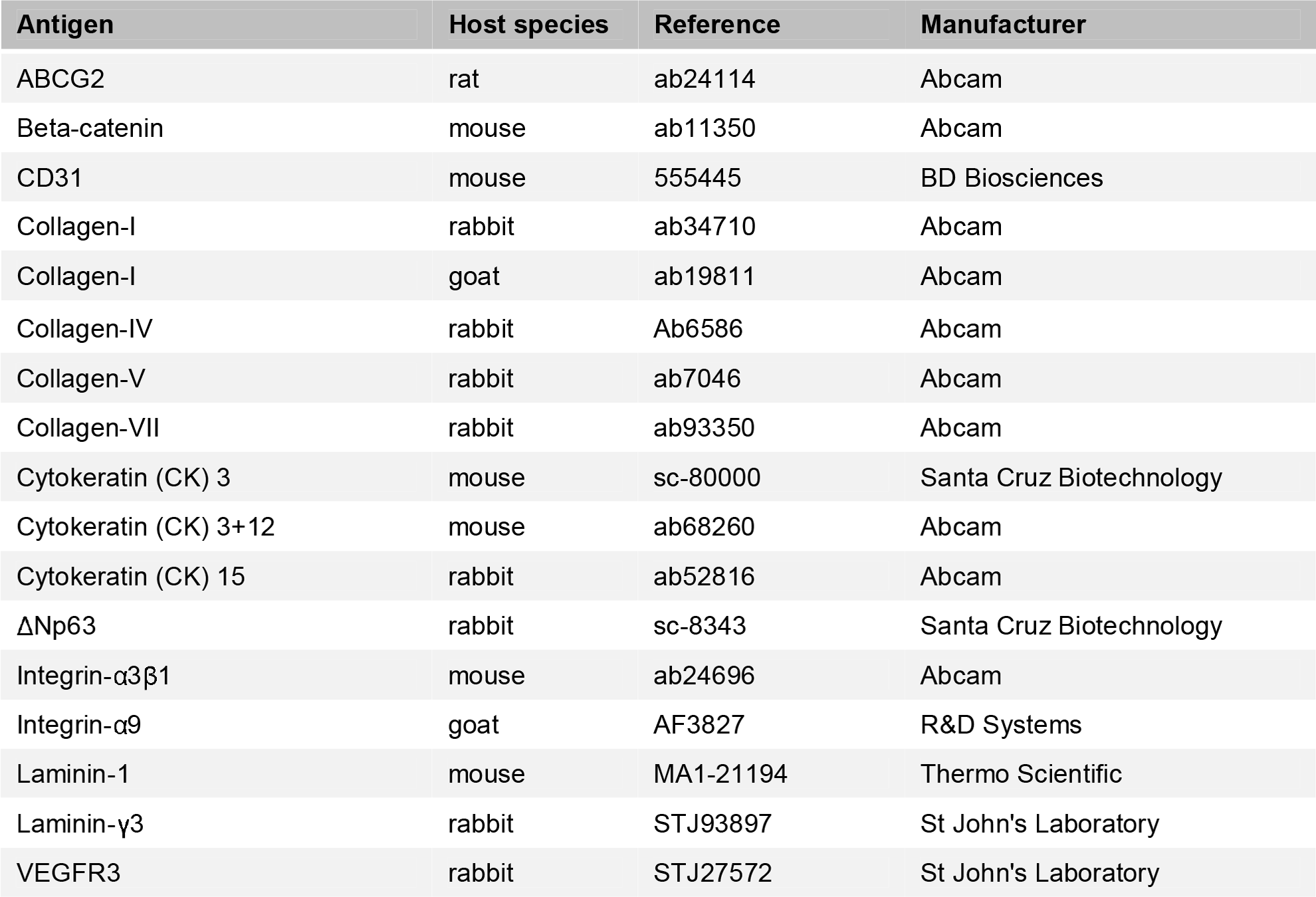
Antibodies used for immunohistochemistry analysis. The manufacturer and corresponding reference numbers are indicated.

### Hydration analysis

Collagen gel weight was measured before treatment (initial weight; Wi) and after treating the entire surface of gels with collagenase for different durations (final weight; Wf). The loss of mass due to collagenase treatment (ΔW) was evaluated for each duration of treatment using the equation ΔW = W_f_ / W_i_ whereas the hydration of treated gels was evaluated by weighting the collagen gels after desiccation using a benchtop Freeze Dry System (Cole-Palmer, IL, USA). Briefly, gels subjected to different treatments were frozen at −54°C to allow water sublimation under controlled pressure for 12 h and their dry weight (W_d_) measured immediately after. Gel hydration was subsequently calculated as the ratio of (W_f_ - W_d_) / W_f_ and expressed as a percentage. The experiment was performed using 10 independent gels for each duration of treatment (*n* = 10).

### Atomic Force Microscopy

Analysis of surface topography of collagen gels treated with collagenase for different durations was performed in static force (contact) mode using an Easyscan 2-controlled FlexAFM atomic force microscope (Nanosurf, Switzerland) equipped with commercial soft contact mode cantilevers (ContAI-G, Budget Sensors, Bulgaria) with a resonant frequency of 13 kHz, and nominal spring constant of 0.2 N/m. Semi-compressed and coated collagen gels were similarly analyzed. Briefly, the different collagen gel samples were mounted onto glass slides that were previously covered with a layer of parafilm to avoid having the very high rigidity of the glass influence the stiffness measurements, and minimize sample displacement and drift. Surface topography was analyzed from three separate regions in each sample, with 512× two-direction lines scanned at 10 µm.s^-1^, 10 nV, and with a P- and I-gain of 1. Surface topography maps were line-fitted and represented in false color with a *z-*scale of 500 nm. The mechanical properties of the collagen gel samples were evaluated by force-distance spectroscopy. The stiffness of the gels was evaluated from 100 force-distance curves acquired at 2 µm.s^-1^ from different positions across each sample, and using SPIP data analysis software (Image Metrology A/S, Denmark) for baseline and hysteresis correction, followed by elastic modulus calculation using the Sneddon model, applicable for soft biological materials (Lin and Horkay, 2008). Elastic modulus was represented in scatter-dot plots or in terms of frequency, in percentage of measured values, within 0.2 and 1.0 MPa bins. All experiments were performed three independent times (*n* = 3).

### Isolation and culture of corneal limbal epithelial stem cells

Corneal ring tissues were kindly provided by Dr Francisco Figueiredo, Royal Victoria Infirmary, in Newcastle-upon-Tyne, UK. Briefly, corneal rings were obtained from cadaverous human donors (ages between 39 and 76; average ± S.D. = 61 ± 12 years; male-female donor ratio of 2:3; no prior history of corneal diseases or ocular trauma) following removal of the central 7 mm for keratectomy. Human corneal rings including the limbus region were dissected into quarters and remaining scleral tissue removed. Tissue quarters were subsequently plated in 6-well plates and incubated for five days in 4 mL of supplemented CnT-7 medium (CellNTec, Switzerland) at 37 °C and 5% CO_2_, to allow the migration and attachment of limbal epithelial cells (hLECs) onto the culture plate surface while maintaining cell proliferation and extended viability. Corneal tissue was then removed from the plates, and attached hLECs allowed to proliferate for an additional week, with culture medium change every two days. Cell monolayers reaching 70-80% confluence were passaged using Accutase cell detachment solution (Thermo Scientific) for 10 min at 37°C, and re-plated at 3 × 10^4^ cells.cm^2^, up to passage 4.

### Proliferation of corneal limbal epithelial cells on collagen gels

Collagen gels treated with collagenase (treated) or PBS (untreated) for 1 h were transferred into Transwell culture inserts (Corning, NY, USA) where they were coated with laminin (1.5 × 10^-6^ g.cm^-2^; Thermo Scientific) for 2 h prior to cell seeding. Human limbal epithelial cells (hLECs) were suspended at 3 × 10^5^ cells per mL of CnT-7 medium and then transferred onto collagen gels (1 mL per insert). All cultures were incubated at 37 °C and 5 % CO_2_ for 2 weeks, with cell culture medium being replaced every 2 days. The AlamarBlue assay was used to assess the proliferation and viability of hLECs grown on treated and untreated collagen gels after 1 week. Briefly, gels were incubated with resazurin reagent (#R7017, Sigma-Aldrich) diluted 1:10 in fresh culture medium and incubated for 4 hours at 37°C, after which 100 µL of culture supernatants were sampled, in triplicate, for fluorescence emission analysis at 590 nm using a Fluoroskan Ascent plate fluorometer (Thermo Scientific). Cell number was calculated by interpolation using a standard curve for fluorescence values of 1, 5, 10, 20, 50, and 100 x 10^4^ cells, with values corresponded to average ± SD of 3 independent experiments (*n* = 3).

### Cell migration assay

Collagen gels previously treated with collagenase (treated) or PBS (untreated) 1 h were laminin-coated, seeded with 5 × 10^5^ hLECs in 1 mL of CnT-7 medium, and incubated at 37°C for 6 hours with the substrate held at an initial 45° tilt to ensure that cells only attached to the lower half-surface of the gel, and with the upper half being above the air-liquid interface. Subsequently, cultures were washed three times with PBS to remove any remaining unattached cells, and incubated at 37°C for 24 h in CnT-7 medium while being imaged at 10 min intervals by time-lapse bright-field microscopy using a Lumascope 500 inverted microscope (Etaluma, CA, USA). Micrographs were binarised using the ImageJ v1.7 software to better determine the position of individual cells in each image frame. Cell speed (µm.h^-1^) was evaluated by determining the position of 100 individual cells during the initial 24 h of migration, and tracing total distance covered by moving cells using the standard parameters of the wrMTrck plugin for ImageJ v1.7. Data was expressed as the average ± S.D. from three independent experiments using cells from independent donors.

### Organ culture of whole human corneas

Collagenase-treated whole human corneas were transferred into Transwell culture inserts after Brillouin spectro-microscopy analysis, washed 3 × 5 min with an excess of sterile PBS to detach and remove existing epithelium, and then seeded with 5 × 10^5^ hLECs in 2 mL of CnT-7 medium. Corneas mock-treated with PBS were used as non-softened controls. For the *ex vivo* alkali burn model, human corneas were similarly processed after treatment with strips of filter paper soaked with 0.5 M of NaOH (alkali) or sterile PBS (control) for 60 s, followed by 3 × 5 min washes with an excess of sterile PBS. A subgroup of alkali burn corneas were also subsequently subjected to collagenase treatment, as described. Organ culture was performed at 37 °C and 5 % CO_2_ in CnT-7 medium for 2 weeks, followed by a 1 week period in SHEM and air-lifting (Jones et al., 2012), with media change every 2 days. The central region of collagenase-treated and control corneas was monitored daily by bright-field microscopy to evaluate re-epithelialization, and then analyzed by confocal immunofluorescence microscopy, as described above. Experiments were performed three independent times (*n* = 3).

### Quantitative RT-PCR analysis

Collagen gels treated with collagenase (treated) or PBS (control) for 60 min were laminin-coated, seeded with 3 × 10^5^ hLECs, and incubated in CnT-7 medium at 37°C for up to 1 week. Cells were then harvested using a flexible cell scraper (Sarstedt, Germany), and their mRNA isolated using standard extraction with Trizol (Thermo Scientific). The assessment of mRNA quality was performed using a Nanodrop 2000 spectrophotometer (Thermo Scientific) to ensure the 260/280 ratio was within the 1.8-2.0 range. Synthesis of cDNA from isolated total mRNA was done using the RT2 First Strand kit (Qiagen, Germany) according to the manufacturer’s protocol, in a TcPlus thermocycler (Techne, UK). The polymerase chain reaction (PCR) was carried out using the default thermal profile of the Eco Real-Time System (Illumina, CA, USA), with the following 40× three-step cycle: 10-second denaturation, 95°C; 30-second annealing, 60°C; and 15-second elongation, 72°C. The transcription levels of *DEFB4* and *KRT3* (differentiation markers), *ABCG2*, *NP63*, *KRT14*, and *NANOG* (LESC markers), and *VEGF*, *IL1A*, *IL1B*, and *IL6* (pro-inflammatory markers) using specific primers (Table 2) were calculated by the comparative threshold cycle (CT) (Eco Software v3.1; Illumina), and normalized to the expression of the *POLR2A* housekeeping gene. Data was represented as gene expression relative to that of hLECs grown on stiffer (control) gels, from 3 independent experiments (*n* = 3).

**Table 2:**
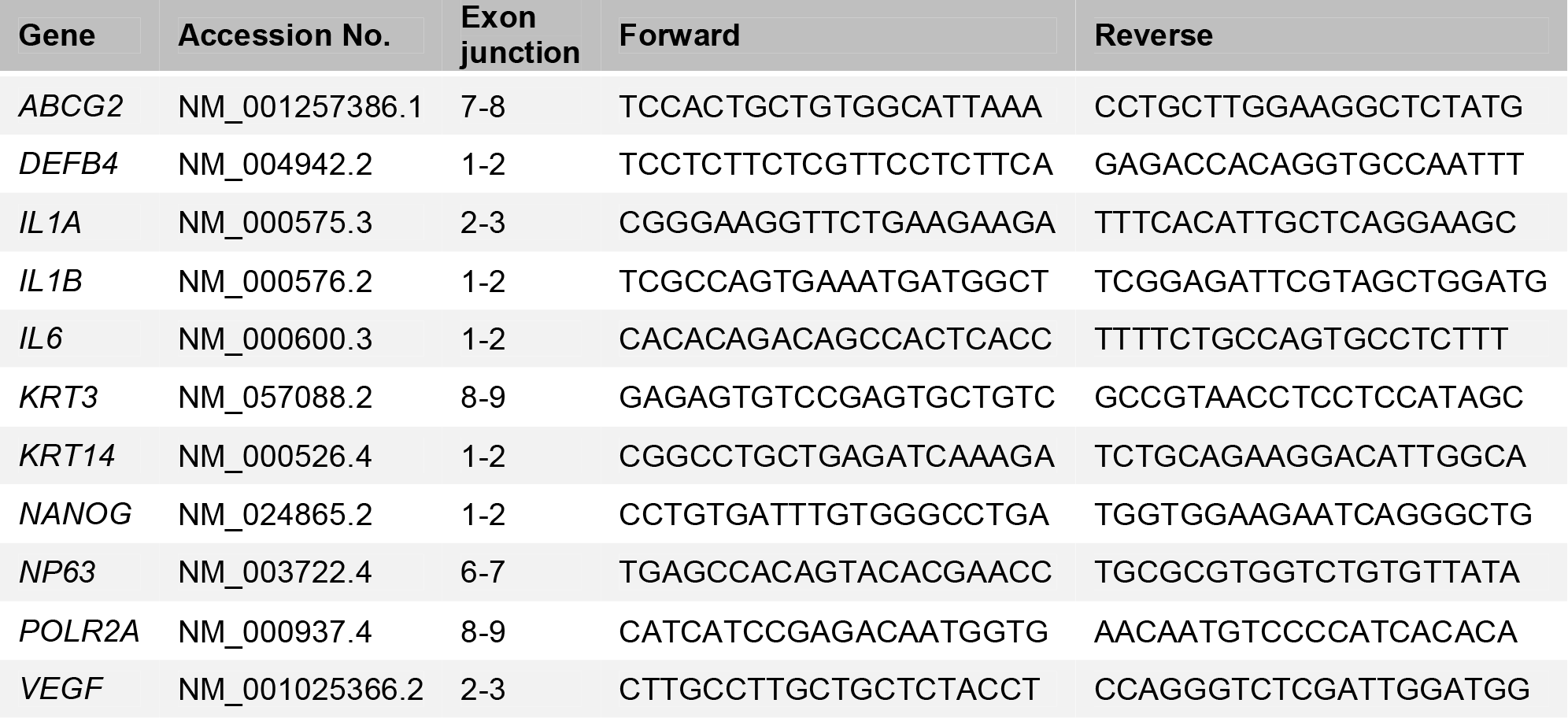
Specific primer pairs used to evaluate gene transcription by quantitative RT-PCR. The corresponding gene accession numbers and location of the amplified sequence within each gene is indicated.

### Quantitative immunofluorescence analysis

Collagen gels treated with collagenase across half their surface or in a ring-shape area for 60 min were laminin-coated, seeded with 3×10^5^ hLECs, and incubated in CnT-7 medium at 37°C for up to 4 weeks. Subsequently, gels were washed twice in PBS, fixed in 4% paraformaldehyde in water for 20 min at room temperature, and washed again with excess PBS. Cells were then permeabilised in PBS containing 0.1% Triton X-100 (PBS-T) for 5 min, blocked with blocking buffer (PBS-T containing 5% BSA) for 1 h, incubated with the rabbit anti-CK3 or anti-CK15 antibodies (Table 1) diluted 1:1000 in blocking buffer for 4 h at room temperature, washed again for 3 × 15 min in blocking buffer, and incubated with IRDye 800CW goat anti-rabbit secondary antibody and CellTag 700 stain diluted 1:1000 in blocking buffer for 2 h at room temperature. Gels were finally washed 3 × 15 min in PBS-T before being imaged by near-infrared quantitative analysis using an Odyssey CLx System (Li-Cor Biosciences, RRID:SCR_014579). Signal specificity was evaluated from gels incubated without either primary or secondary antibodies. The intensity of signal corresponding to cellular CK3 and CK15 detection was determined for the collagenase-treated and untreated areas of the gels, and normalized to the number of cells present on the corresponding areas, determined by the CellTag stain signal. Marker expression was also compared to that from cells grown on tissue culture plastic for corresponding periods of time. Data was represented as protein expression relative to the average expression of untreated (control) areas of the gels from three independent experiments using cells from independent donors.

### Enzyme-linked immunosorbent assay

Collagen gels treated with collagenase (treated) or PBS (control) for 60 min were laminin-coated, seeded with 3×10^5^ hLECs, and incubated in CnT-7 medium at 37°C for 3 days. Subsequently, culture supernatants were collected on ice, centrifuged at 10,000 ×*g* for 10 min at 4°C, sampled, and analyzed for human IL-6 expression using the Quantikine enzyme-linked immunosorbent assay (ELISA) kit (R&D Systems, RRID:SCR_006140) according to the manufacturer’s instructions. The expression of IL-6 was then normalized for the amount of total protein present in the supernatants, and expressed as pg.mL^-1^.µg^-1^ of total protein. Both conditions were assayed in triplicate.

### Collagenase treatment of rabbit corneas

Animal experiments were performed following the appropriate guidelines and after ethical approval from the Institutional Animal Care and Use Committee of the Harry S. Truman Memorial Veterans’ Hospital and the University of Missouri, Columbia, USA. To test the effect of collagenase on the central cornea, two to three month old New Zealand White female rabbits (Covance Research Products, RRID:SCR_001224) weighing 2.5 to 3.0 kg were anesthetized and divided into two groups of six animals: one subjected to collagenase treatment in their intact (right eye) and debrided (left-eye) corneas, previously marked in the central region using an 8 mm Ø trephine and debrided of the epithelium; and a second group subjected to mock-treatment. The effect of collagenase on chemically-burned limbus tissue was tested using the right eyes of six rabbits subjected to the localized application of 0.5 M NaOH for 25 s (alkali burn) in the temporal side of the limbus, with a sub-group of three animals subsequently receiving collagenase treatment on the burned limbus (treated) (see Supplementary Figure 9). The softening treatment was performed based on the collagenase activity optimized previously *in vitro* and *ex vivo*. Briefly, collagenase was dissolved in 100 µL of pre-warmed 1:19 Dextran-PBS sterile solution (0.2 g.L^-1^; 4×10^4^ activity units.L^-1^), soaked into an Ultracell Corneal Light Shield PVA sponge (Beaver-Visitec, MA, USA), and then applied to the central or limbus region of the cornea for 15 min (2 units.cm^-2^.h^-1^ of total collagenase activity). Mock-treatments were performed by applying Dextran-PBS-soaked sponges to the cornea. Corneas were then washed thoroughly by excess saline and evaluated 1 and 5 days (central cornea softening assay) or 0, 2, and 7 days (alkali-burned limbus assay) post-intervention by intraocular pressure (IOP) measurements, unbiased clinical eye examination, and stereo- and slit lamp-microscopy. At each time point, six rabbits (3 collagenase-treated, 3 control animals) were sacrificed after examination, and the whole corneas excised, embedded in Optical Cutting Temperature (OCT) medium, and stored at −80°C until processed for immunohistochemistry and fluorescence confocal microscopy analysis, as described above.

### Statistical analysis

Error bars represent the standard deviation (S.D.) of the mean, analyzed *a priori* for homogeneity of variance. Replicates from each independent experiment were confirmed to follow a Gaussian distribution, and differences between groups were determined using one- or two-way analysis of variance (ANOVA) followed by Bonferroni’s multiple comparison *post hoc* test. Significance between groups was established for *p* < 0.05, 0.01, and 0.001, and with a 95% confidence interval. The average ± S.D. Brillouin frequency shift was calculated using 100 points from discrete corneal tissues (epithelium, sub-epithelial lamina, anterior stroma, sub-limbal conjunctiva) from three independent donors (*n* = 3), and represented as a scatter plot to illustrate intra-tissue variations. The average ± S.D. frequency distribution of elastic modulus of both treated and untreated gels was tested for correlation, and fit to Gaussian curves using a non-linear least-squares regression model, which were then analyzed for independence using the *Chi*-square test.

